# SMALL ALPHAHERPESVIRUS LATENCY-ASSOCIATED PROMOTERS DRIVE EFFICIENT AND LONG-TERM TRANSGENE EXPRESSION IN THE CENTRAL NERVOUS SYSTEM

**DOI:** 10.1101/2019.12.31.891903

**Authors:** Carola J. Maturana, Jessica L. Verpeut, Thomas J. Pisano, Zahra M. Dhanerawala, Andrew Esteves, Lynn W. Enquist, Esteban A. Engel

## Abstract

Recombinant adeno-associated viral vectors (rAAV) are used as gene therapy vectors to treat central nervous system (CNS) diseases. Despite their safety and broad tropism, important issues need to be corrected such as the limited payload capacity and the lack of small gene promoters providing long-term, pan-neuronal transgene expression in the CNS. Commonly used gene promoters are relatively large and can be repressed a few months after CNS transduction, risking the long-term performance of single-dose gene therapy applications. We used a whole-CNS screening approach based on systemic delivery of AAV-PHP.eB, iDisco+ tissue-clearing and light-sheet microscopy, to identify three small latency-associated promoters (LAP) from the herpesvirus pseudorabies virus (PRV). These promoters are LAP1 (404bp), LAP2 (498bp) and LAP1_2 (880bp). They drive chronic transcription of the virus encoded latency-associated transcript (LAT) during productive and latent phases of PRV infection. We observed stable, pan-neuronal transgene transcription and translation from AAV-LAP in the CNS for six months post AAV transduction. In several CNS areas, the number of cells expressing the transgene was higher for LAP2 than the large conventional EF1α promoter (1264bp). Our data suggests that the LAP are suitable candidates for viral vector-based CNS gene therapies requiring chronic transgene expression after one-time viral-vector administration.

## Introduction

Recent improvements to recombinant adeno-associated viruses (AAVs), including capsid engineering and novel gene promoters to optimize transgene expression have substantially improved gene therapy applications^1–3^. AAV vectors are widely used in neuroscience and clinical applications given their safety, serotype-dependent broad tropism and transduction efficiency^1, 4–6^. AAV-9 variant PHP.eB^7^, with an enhanced ability to permeate the mouse blood-brain barrier (BBB) and broadly transduce CNS neurons both in the brain and spinal cord after peripheral vascular administration, is one example of recent capsid improvements^8^. A major limitation of recombinant AAVs is their small capsid with limited payload capacity of only ~ 4.9 kb^9^. Accordingly, the discovery of short promoter sequences that sustain strong and long-lived transcription is paramount to expand the transgene payload and achieve chronic therapeutic effect with one viral dose.

Several strong promoters such as neuron-specific enolase (NSE, 1800 bp)^10, 11^, calcium/calmodulin-dependent protein kinase II alpha (CaMKIIa, 1300 bp)^12^ and human elongation factor 1 alpha (EF1α, 1264 bp)^12, 13^ have been used in systemic AAV delivery^6^. However, the considerable size of these promoter sequences limits the use of large therapeutic transgenes or multiple small transgenes. Moreover, short promoters such as the human cytomegalovirus immediate-early enhancer and promoter (CMV, 600 bp)^14^ or truncated versions of the human synapsin promoter (hSyn, 468 bp)^15^, are considerably weaker to drive gene transcription and expression, and in some cases, are completely repressed or inactivated only weeks after delivery ^13, 15–18^. Similarly, small ubiquitous promoters like beta glucuronidase (GUSB, 378 bp)^19^ or ubiquitin C (UBC, 403 bp)^13, 20^ have shown weak transcription levels.

Here, we describe and validate three alphaherpesvirus latency-associated promoters (LAP), called LAP1 (498 bp), LAP2 (404 bp) and LAP 1_2 (880 bp) obtained from the genome of the herpesvirus pseudorabies virus (PRV). The *Alphaherpesvirinae* subfamily of the family *Herpesviridae*, includes bovine herpes virus-1 (BHV-1), varicella-zoster virus (VZV), herpes simplex virus (HSV) and PRV. These viruses share genome organization and establish latent infections in sensory ganglia of different mammalian hosts^21–23^. Several studies have established the role of LAP in promoting long-term expression, since LAP can chronically drive transcription of the latency-associated transcript (LAT), even under highly repressible and adverse conditions^24–30^. The LAP region of PRV encompasses two independent promoters, LAP1 and LAP2^27, 31–33^. PRV LAP1 contains two GC boxes and three CAAT boxes upstream of the first TATA box. PRV LAP2 contains two GC boxes before the second TATA box^27, 32, 34^. It has been proposed that the binding of different transcription factors (TF) to consensus promoter elements present in LAP may facilitate escape from nucleosome silencing during the latent infection^33, 35–38^. In transgenic mouse lines, PRV LAP promoted transcription is neuron-specific in the absence of PRV infection^39^. However, in transient expression assays, PRV LAP1 and LAP2 promoted transcription both in cultured neuronal as well as non-neuronal cells^31, 34^. Furthermore, the activity of tandem LAP1 and LAP2 sequences is significantly increased compared to LAP1 or LAP2 alone^31^.

In the present study, we tested PRV LAP promoters in AAV vectors for *in vivo* whole-CNS transduction. We found that PRV LAP1, LAP2 and tandem LAP1_2 promoters are suitable for systemic, less invasive, pan-neuronal gene delivery applications that may require stable, chronic transgene expression after a single administration.

## Results

### Small PRV LAP variants can drive transgene expression in neurons independently of herpesvirus infection

The PRV LAP region includes at least two promoter regions defined here as LAP1 and LAP2 (Figure 1A). In the PRV genome, LAP1 and LAP2 are present in tandem as PRV LAP1_2. These sequences alone or combined are capable of efficient expression of reporter transgenes in primary sympathetic neurons when used in AAV vectors without PRV infection (Figure 1 C and 1D). We analyzed the LAP nucleotide sequences to identify putative regulatory elements using Jasper^40, 41^, RSAT^42^ and CTCFBSDB 2.0^43^ software. We identified three cyclic AMP response element-binding protein (CREB) located upstream of the LAP1 TATA box and one upstream of LAP2 TATA box. Moreover, two CTCF motifs (CCCTC - binding factor) were detected upstream of LAP1 TATA box and one downstream of LAP2 TATA box. We identified downstream promoter elements (DPE) in LAP2, including CG boxes and four signal transducer and activator of transcription 1 (STAT1) sites. Additionally, there were lineage-determining TF^44^, such as SRY-Box 10 (SOX10) and oligodendrocyte transcription factor 2 (Olig2), upstream of the LAP1 TATA box and LAP2 TATA box, respectively (Figure 1A).

**Figure 1.**
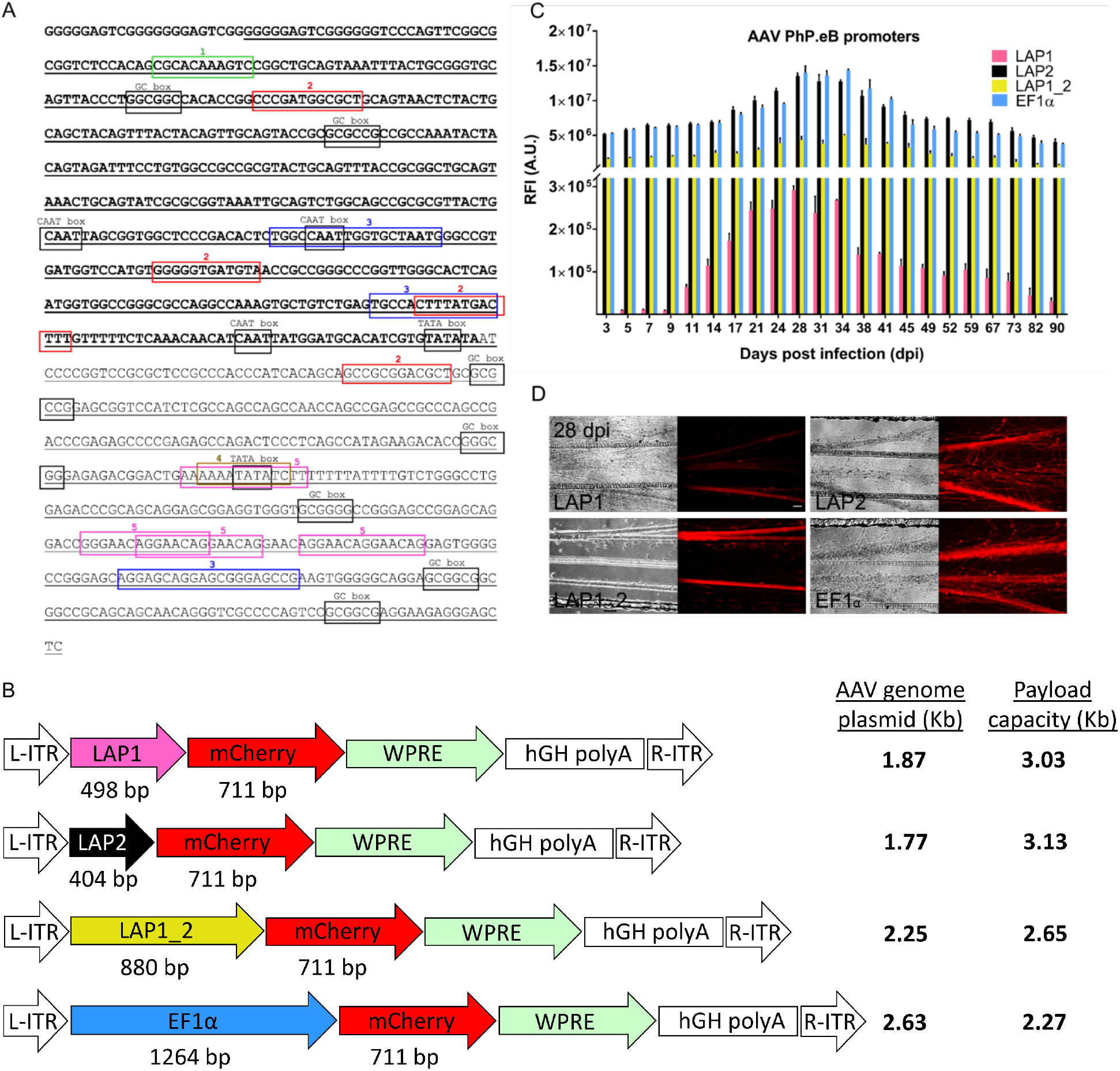
Characterization of PRV latency-associated transcript promoter (LAP). (A) The complete nucleotide sequence of PRV LAP of 902 bp, and sub-regions LAP1 (bold and underlined) of 498 bp, LAP2 (underlined) of 404 bp, and LAP1_2 of 880 bp are depicted. LAP1_2 includes most of LAP sequence, except for the first 22 nt of LAP1. Black boxes depict consensus sequences for TF including, GC box: specificity protein 1 and 3 (Sp1 and Sp3); CAAT box: nuclear factor Y (NF-Y); TATA box: TATA-binding protein (TBP). Colored boxes indicate the coordinates for TF binding motif sites: 1, green: SRY-Box 10 (SOX10); 2, red: cAMP response element-binding protein (CREB); 3, blue: CCCTC-binding factor (CTCF); 4, brown: oligodendrocyte transcription factor 2 (Olig2); 5, pink: signal transducer and activator of transcription (STAT1). (B) Plasmid maps of four AAVs designed to transcribe mCherry fluorescent reporter from promoters LAP1, LAP2, LAP1_2 and EF1α. WPRE of 609 bp is a woodchuck hepatitis virus posttranscriptional enhancer element. All AAVs contain a 479 bp human growth hormone (hGH polyA) poly adenylation sequence, and flanking AAV2 inverted terminal repeats (ITRs) of 141 bp each. Vectors were packaged into the AAV-PhP.eB serotype capsid. The total size of the enhancer-promoter elements and promoter sequence is AAV-LAP1 (1.87 Kb), AAV-LAP2 (1.77 Kb), AAV-LAP1_2 (2.25 Kb) and AAV-EF1α (2.63 Kb) respectively. (C) All four AAVs were used to transduce primary cultures of rat SCG neurons to quantify mCherry expression over a 90 day time-lapse. The relative fluorescence intensity of mCherry expression was measured at 3, 5, 7, 9, 11, 14, 17, 21, 24, 28, 31, 34, 38, 41, 45, 49, 52, 59, 67, 73, 82, and 90 days post infection (dpi) with 3 × 10^11^ vg. Data are represented as mean ± SEM; n = 3 SCG culture dishes per group. (D) AAV-driven mCherry expression in SCG neurons is shown at 28 dpi with LAP1-mCherry, LAP2-mCherry, LAP1_2-mCherry and EF1α-mCherry. Scale bar = 500 μm.

Four AAV recombinants were packaged into serotype PHP.eB capsids by standard methods. We constructed three promoters: LAP1 (498 bp), LAP2 (404 bp) and LAP 1_2 (880 bp). We used the ubiquitous EF-1α promoter (1264 bp) as a positive control for transgene expression. All four AAV recombinants expressed the fluorescent reporter mCherry (Figure 1B). To verify the *in vitro* performance of each promoter, we transduced rat primary superior cervical ganglia (SCG) neuronal cultures with 3 × 10^11^ genomes of each AAV and quantified the relative mCherry fluorescence intensity (RFI) over a 90-day period. For neurons transduced with AAV-LAP1, mCherry expression increased abruptly at 11 days-post-infection (dpi) (6.42^4^ RFI) but to a lower level when compared with AAV-LAP2, AAV-LAP1_2 and AAV-EF1α. AAV-LAP2 and AAV-EF1α expression increased ~125-fold more at 17 dpi (8.70 × 10^6^; 8.01 × 10^6^ RFI respectively) (Figure 1C). The highest level of expression by all four recombinants was at 28 dpi with LAP1 (2.9 × 10^5^ RFI); LAP2 (1.36 × 10^7^ RFI); LAP1_2 (4.36 × 10^6^ RFI) and EF1α (1.40 × 10^7^ RFI) (Figure 1C and 1D). LAP2 and EF1α had the highest mCherry RFI (LAP2=EF1α>LAP1_2>LAP1). Between 38 dpi and 90 dpi, all four AAV recombinants showed a subtle but sustained RFI decrease (Figure 1C), most likely due to the senescence of primary SCG neurons after more than 100 days in culture. Importantly, all three AAV-LAP recombinants showed mCherry transcription in primary neurons for 90 days in the context of AAV transduction and in the absence of PRV infection.

### Whole-CNS screening reveals pan-neuronal AAV-LAP transgene expression after six months

We used AAV serotype PHP.eB for our promoter screening assay given the enhanced capacity to cross the BBB and transduce C57/BL6 mice CNS after systemic, intravascular delivery^7^. AAV-LAP1-mCherry, AAV-LAP2-mCherry, AAV-LAP1_2-mCherry and AAV-EF1α-mCherry were delivered by unilateral retro-orbital venous sinus injection of 4 × 10^11^ viral genomes/mouse (vg/mouse). The brains and spinal cords were harvested 30 dpi and 190 dpi, as described in Figure 2A, to quantify mCherry transcription and translation. LAP-mCherry expression in the whole intact brain was determined after tissue was cleared and immunostained with iDISCO+ (immunolabeling-enabled three-dimensional imaging of solvent-cleared organs) by light-sheet microscopy and volumetric registration (Supplementary Movies S2, S3, S4, S5)^45, 46^. All four gene promoters showed stable mCherry expression at both 30 and 190 dpi. The density (number of mCherry positive cells per mm^3^ of brain tissue) of LAP2 was higher than that of LAP1 and LAP1_2, and had no significant differences to EF1α (p < 0.05) in different areas of cortex: primary motor, secondary motor, primary somatosensory and supplemental somatosensory (Figures 2B – 2E respectively); hippocampal formation (Figure 2G), pallidum (Figure 2I), hypothalamus (Figure 2K) and olfactory areas (Figure 2P). In cerebellum, LAP2 showed significatively higher mCherry density than LAP1, LAP1_2 and EF1α (Figure 2O). Furthermore, in striatum (Figure 2H); thalamus (Figure 2J); midbrain, motor and sensory areas (Figure 2L and 2M), and hindbrain (Figure 2N), LAP2 and LAP1_2 showed significantly higher density than EF1α (p < 0.05). Note that the LAP2 nucleotide sequence is 68% shorter than that of EF1α yet outperforms EF1α density in several brain areas. To further validate LAP transgene expression in the CNS, we assessed mCherry protein expression by immunohistochemistry (IHC) in brain sagittal sections at 30 and 190 dpi (Figure 3A - 3D). Analysis by confocal microscopy, showed abundant mCherry staining throughout the cortical somatosensory area (Figure 3E1 - 3E4), dentate gyrus in the hippocampal formation (Figure 3F1 - 3F4), caudoputamen in the striatum (Figure 3G1 - 3G4), and cerebellar cortex (Figure 3H1 - 3H4) at 30 dpi. Importantly, mCherry expression was stable for all three LAP variants at 190 dpi and similar to that of the large promoter EF1α (Figure 3E5 - 3E8, 3F5 - 3F8, 3G5 - 3G8 and 3H5 - 3H8). Next, we quantified mCherry expression at 30 and 190 dpi. The mCherry RFI was similar for all AAV promoters with no significant differences (p < 0.05) (Figure 3I1 - 3I4). We subsequently quantified the number of mCherry positive cells per pixels^2^ 190 dpi. In cortex, the number of LAP2-mCherry expressing cells was higher than those observed for LAP1-mCherry and LAP1_2-mCherry [LAP2: 297 ± 19.82 vs. LAP1: 149 ± 5.61 vs LAP1_2: 168 ± 9.22 (n = 6, p <0.001) (Figure 3J1). In dentate gyrus, striatum and cerebellum the number of mCherry positive cells was similar for all LAP variants and EF1α (Figure 3J2 - 3J4). We conclude that all AAV-LAP variants promote mCherry expression in the brain, further demonstrating that a single administration of AAV-LAP recombinants is sufficient to drive long term, pan-neuronal transgene expression in the mouse CNS.

**Figure 2.**
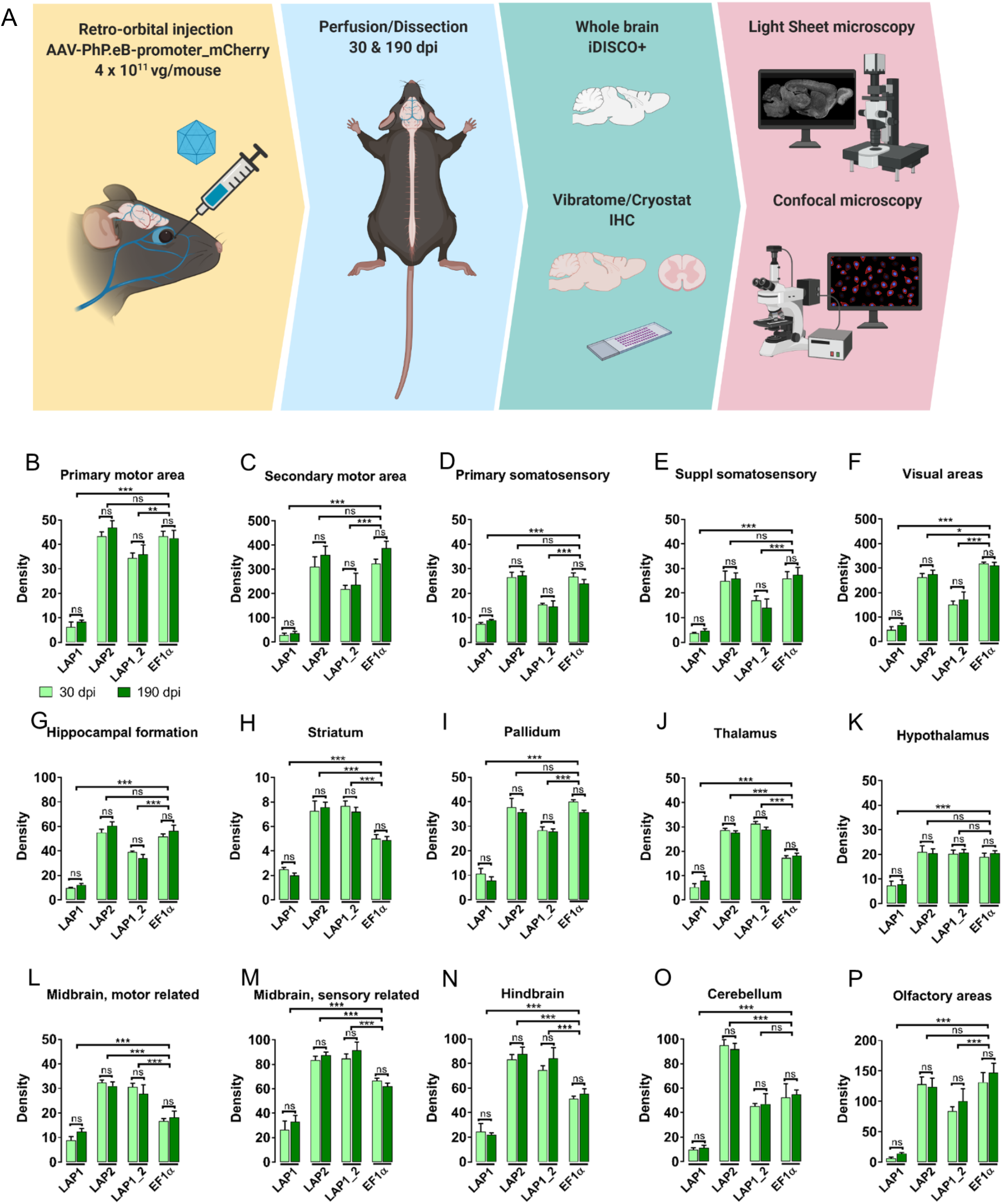
Whole-brain volumetric registration of AAV-infected animals show stable and long-term LAP-mediated transgene expression. (A) Schematic of the systemic route of AAV administration and subsequent CNS tissue processing. Intravenous administration of AAV vectors was performed by unilateral injection into the mice retro-orbital sinus (4 × 10^11^ vg/mouse). Brain and spinal cord were collected at 30 and 190 dpi. Brain right hemispheres were processed for iDISCO+ tissue clearing and subsequent light-sheet microscopy analysis. The left hemispheres were sagittally sectioned at 50 μm for subsequent IHC and confocal microscopy analyses. Spinal cords were transversally sliced at 20 μm for subsequent confocal microscopy analyses. (B-P) Quantification of the density of mCherry positive cells per mm^3^ across different brain regions in iDISCO+ tissue cleared samples at 30 and 190 dpi. Cortex area: (B) primary motor area, (C) secondary motor area, (D) primary somatosensory area, (E) supplemental somatosensory area, (F) visual areas; (G) hippocampal formation; (H) striatum; (I) pallidum; (J) thalamus; (K) hypothalamus; midbrain area: (L) motor related, (M) sensory related; (N) hindbrain; (O) cerebellum; (P) olfactory areas. Data was normalized to a vehicle-injected control animal. Data are represented as mean ± SEM; n = 2 animals per group. Between five and ten 500 um volumes per region were analyzed for each animal. Data was normalized to a vehicle-injected control animal. Significance was determined with Student’s t-test (if only two groups were compared) or analysis of variance one-way (ANOVA) followed by Bonferroni post hoc test (if more than two groups were compared). A p value <0.05 was considered to be statically significant (*p < 0.033; **p < 0.002; ***p < 0.001).

**Figure 3.**
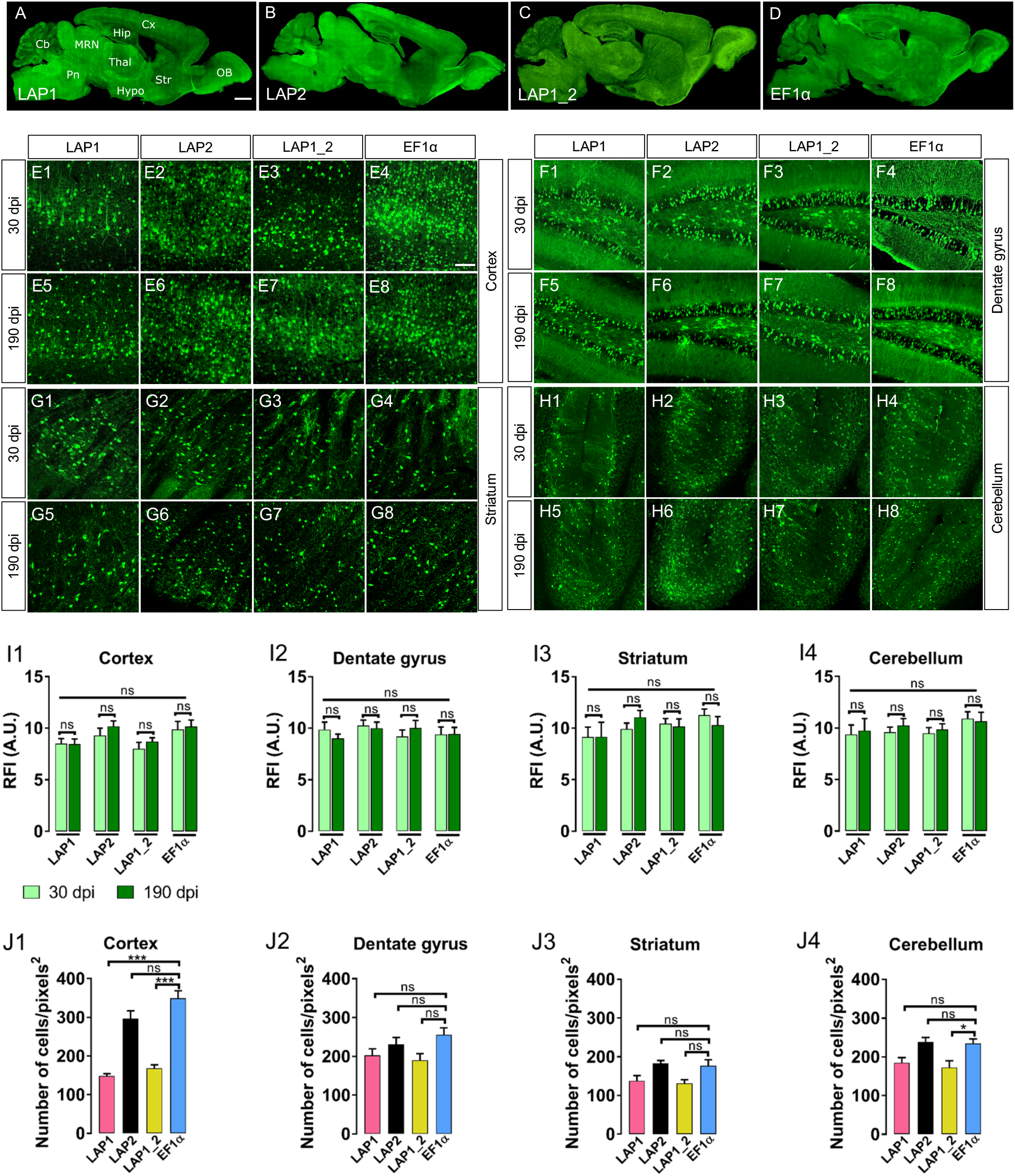
All three AAV-LAP-mCherry variants exhibit widespread and long-term transgene expression throughput the brain after retro-orbital injection. Representative immunofluorescence images of sagittal sections showing whole-brain distribution of anti-mCherry staining are shown in green for (A) AAV-LAP1, (B) AAV-LAP2, (C) AAV-LAP1_2, and (D) AAV-EF1α at 190 dpi. Cx, cortex; Hip, hippocampus; MRN, midbrain reticular nucleus; Cb, cerebellum; Thal, thalamus; Pn, pons; Hypo, hypothalamus; Str, striatum; OB, olfactory bulb. Scale bar = 1mm. Representative confocal images show anti-mCherry signal (green) for AAV-LAP1 (E1, E5) in cortex, (F1, F5) dentate gyrus, (G1, G5) striatum and (H1, H5) cerebellum at 30 and 190 dpi, respectively. AAV-LAP2 (E2, E6) in cortex, (F2, F6) in the dentate gyrus, (G2, G6) in the striatum and (H2, H6) in cerebellum at 30 and 190 dpi, respectively. AAV-LAP1_2 (E3, E7) in cortex, (F3, F7) in the dentate gyrus, (G3, G7) in the striatum and (H3, H7) in cerebellum at 30 and 190 dpi, respectively. AAV-EF1α (E4, E8) in cortex, (F4, F8) in the dentate gyrus, (G4, G8) in the striatum and (H4, H8) in cerebellum at 30 and 190 dpi, respectively. All images are stack confocal sections. Scale bar = 100 μm. Quantification of the indirect fluorescence intensity of anti-mCherry signal driven by AAV-LAP variants and AAV-EF1α at 30 and 190 dpi is shown in (I1) cortex, (I2) dentate gyrus, (I3) striatum, (I4) cerebellum. Quantification of the number of cells expressing mCherry signal per pixels^2^ by IHC at 190 dpi is shown in (J1) cortex, (J2) dentate gyrus, (J3) striatum and (J4) cerebellum. Data are represented as mean ± SEM; n = 2 (six tissue sections were analyzed for each animal). For A, data was normalized to a vehicle-injected control animal. Significance was determined with Student’s t-test (if only two groups were compared) or analysis of variance one-way (ANOVA) followed by Bonferroni post hoc test (if more than two groups were compared). A p value < 0.05 was considered to be statically significant (*p < 0.033; **p < 0.002; ***p < 0.001).

### The small LAP2 promoter variant drives strong and stable pan-neuronal transgene transcription and translation after systemic AAV administration

We compared the efficacy of mCherry expression under the control of different PRV LAP variants We observed abundant signal in the cortex (Figure 4A1 - 4A8, 4E1), dentate gyrus (Figure 4B1 - 4B8, 4E2), striatum (Figure 4C1 - 4C8, 4E3) and cerebellum (Figure 4D1 - 4D8, 4E4) at 30 and 190 dpi. Moreover, the AAV-LAP2 RFI was stable and similar to AAV-EF1α both at 30 and 190 dpi (p < 0.05) (Figure 4E1 - 4E4, Table 1). Although AAV-LAP1 and AAV-LAP1_2 mCherry RFI levels were stable and not significantly different at 30 and 190 days in cortex, dentate gyrus, striatum and cerebellum (Figure 4E1 – 4E4, Table 1), both promoters showed significantly less transgene expression (RFI) compared to LAP2 and EF1α.

**Table 1.** mCherry expression in the mouse brain with different AAV-PHP.eB-promoters.

| Promoter | Brain region | Direct mCherry expression |  | Indirect IHC<sup>a</sup> mCherry expression |  |
| --- | --- | --- | --- | --- | --- |
|  |  | 30 dpi<sup>b</sup> | 190 dpi | 30 dpi | 190 dpi |
|  |  | Mean (RFI)<sup>c</sup> ± SEM | Mean (RFI) ± SEM | Mean (RFI) ± SEM | Mean (RFI) ± SEM |
| **LAP1** | Cortex | 1.05 ± 0.12 | 1.35 ± 0.14 | 8.51 ± 0.47 | 8.46 ± 0.49 |
|  | Dentate gyrus | 1.36 ± 0.11 | 0.99 ± 0.04 | 9.87 ± 0.72 | 9.00 ± 0.44 |
|  | Striatum | 1.18 ± 0.13 | 1.41 ± 0.18 | 9.14 ± 0.97 | 9.17 ± 1.41 |
|  | Cerebellum | 1.31 ± 0.17 | 1.48 ± 0.15 | 9.38 ± 0.92 | 9.75 ± 1.16 |
| **LAP2** | Cortex | 9.66 ± 0.41 | 9.39 ± 0.69 | 9.30 ± 0.70 | 10.18 ± 0.51 |
|  | Dentate gyrus | 8.61 ± 0.63 | 9.32 ± 0.63 | 10.26 ± 0.52 | 10.01 ± 0.59 |
|  | Striatum | 9.91 ± 0.51 | 10.61 ± 0.65 | 9.92 ± 0.57 | 11.06 ± 0.66 |
|  | Cerebellum | 10.66 ± 0.67 | 10.34 ± 0.84 | 9.59 ± 0.50 | 10.26 ± 0.65 |
| **LAP1_2** | Cortex | 5.18 ± 0.43 | 5.36 ± 0.39 | 8.01 ± 0.61 | 8.70 ± 0.36 |
|  | Dentate gyrus | 3.92 ± 0.70 | 4.17 ± 0.62 | 9.20 ± 0.64 | 10.02 ± 0.72 |
|  | Striatum | 4.36 ± 0.84 | 4.67 ± 0.75 | 10.45 ± 0.48 | 10.17 ± 0.75 |
|  | Cerebellum | 4.76 ± 0.54 | 5.42 ± 0.27 | 9.48 ± 0.56 | 9.86 ± 0.54 |
| **EF1α** | Cortex | 8.99 ± 0.62 | 9.05 ± 0.73 | 9.87 ± 0.77 | 10.16 ± 0.62 |
|  | Dentate gyrus | 9.16 ± 0.57 | 9.16 ± 0.81 | 9.40 ± 0.72 | 9.45 ± 0.64 |
|  | Striatum | 9.96 ± 0.59 | 10.93 ± 0.67 | 11.26 ± 0.61 | 10.29 ± 0.85 |
|  | Cerebellum | 10.72 ± 0.49 | 11.06 ± 0.84 | 10.92 ± 0.66 | 10.65 ± 0.89 |

<sup>a</sup>IHC: immunohistochemistry
<sup>b</sup>dpi: days post AAV injection
<sup>c</sup>RFI: relative fluorescence intensity

**Figure 4.**
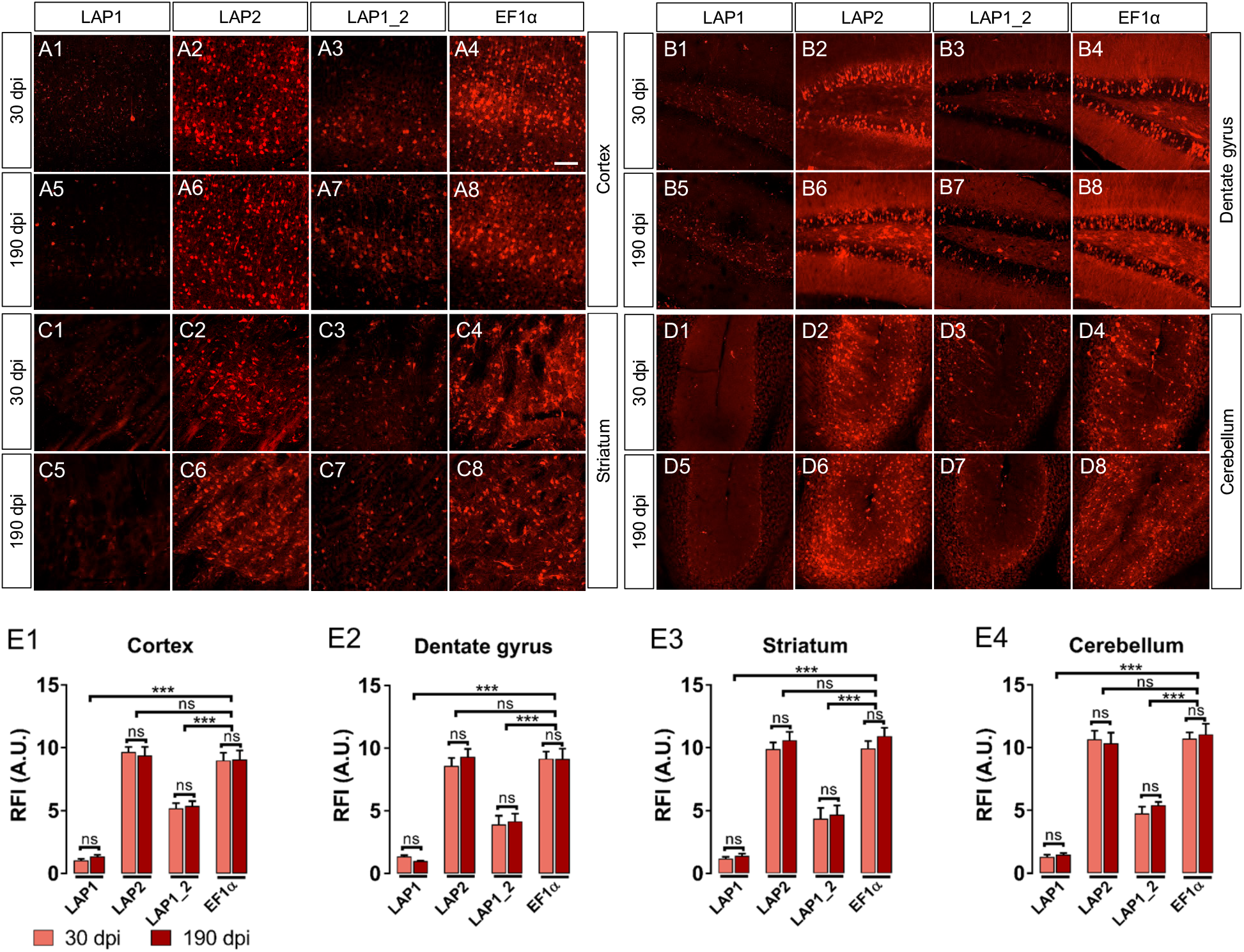
LAP2 drives stable and long-term transgene expression in the brain. Representative confocal images show native mCherry fluorescence (red) for AAV-LAP1 (A1, A5) in cortex, (B1, B5) dentate gyrus, (C1, C5) striatum and (D1, D5) cerebellum at 30 and 190 dpi, respectively. AAV-LAP2 (A2, A6) in cortex, (B2, B6) in the dentate gyrus, (C2, C6) in the striatum and (D2, D6) in cerebellum at 30 and 190 dpi, respectively. AAV-LAP1_2 (A3, A7) in cortex, (B3, B7) in the dentate gyrus, (C3, C7) in the striatum and (D3, D7) in cerebellum at 30 and 190 dpi, respectively. AAV-EF1α (A4, A8) in cortex, (B4, B8) in the dentate gyrus, (C4, C8) in the striatum and (D4, D8) in cerebellum at 30 and 190 dpi, respectively. All images are stack confocal sections. Scale bar = 100 μm. Quantification of the direct fluorescence intensity (RFI) of native mCherry signal driven by AAV-LAP variants and AAV-EF1α at 30 and 190 dpi is shown in (E1) cortex, (E2) dentate gyrus, (E3) striatum, (E4) cerebellum. Data are represented as mean ± SEM; n = 2 (six tissue sections were analyzed for each animal) and was normalized to a vehicle-injected control animal. Significance was determined with Student’s t-test (if only two groups were compared) or analysis of variance one-way (ANOVA) followed by Bonferroni post hoc test (if more than two groups were compared). A p value < 0.05 was considered to be statically significant (*p < 0.033; **p < 0.002; ***p < 0.001).

Since mRNA half-life is typically shorter than that of the translated protein^47^, we measured mCherry transcripts in AAV-LAP2 and AAV-EF1α transduced brains 190 dpi with a mCherry-specific riboprobe. Fluorescent *in situ* hybridization (FISH) showed abundant AAV-LAP2 mCherry RNA in cortex, dentate gyrus, striatum, cerebellum and olfactory bulb (Figure 5), further confirming that PRV LAP2 can drive chronic and robust transgene transcription in the CNS.

**Figure 5.**
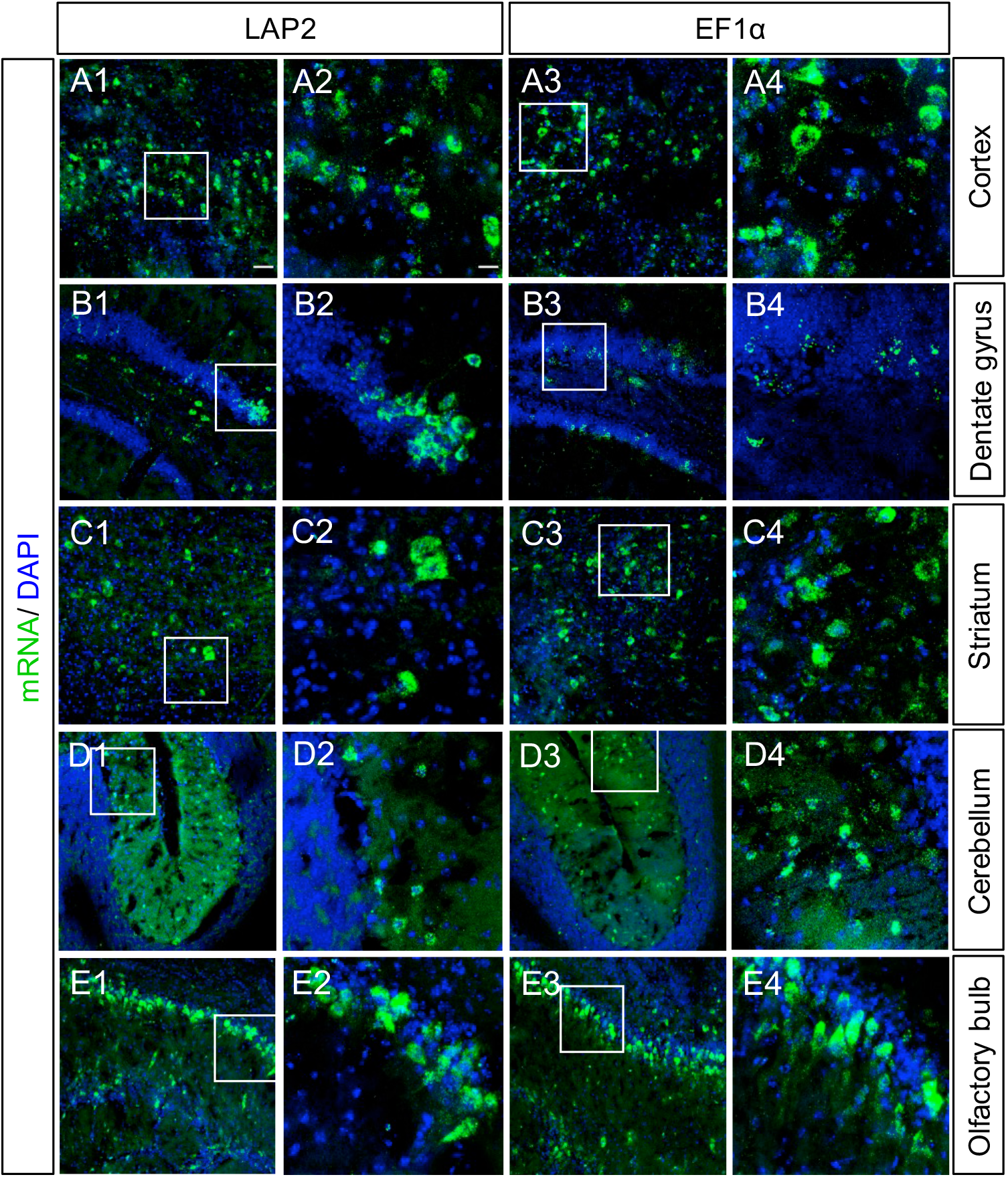
AAV-LAP2 drives long-term transgene transcription in the CNS. The presence of mCherry mRNA was verified by FISH in brain tissue at 190 dpi using a riboprobe specific to mCherry (green). Nuclei were counterstained with DAPI (blue). Brain sagittal 20 μm slices are shown for AAV-LAP2 and AAV-EF1α (A) in cortex, (B) dentate gyrus, (C) striatum, (D) cerebellum and (E) olfactory bulb respectively. (A2, A4, B2, B4, C2, C4, D2, D4, E2, E4) higher magnification images (white square) of each region are shown in A1, A3, B1, B3, C1, C3, D1, D3, E1, E3 respectively. Images are stack confocal sections, scale bars = 50 μm and 100 μm.

### AAV-LAP transgene expression in the brain is predominant in neurons but not in glial cells

The tropism and specificity of AAV transduction and subsequent transgene expression depends on the AAV serotype^4, 6^ and the gene promoter^8, 48, 49^. To characterize which cell-types showed AAV-LAP-mCherry expression after systemic AAV-PHP.eB delivery, we performed co-immunostaining of mCherry protein with markers for neurons (NeuN), oligodencrocytes (Olig2), microglia (Iba1) and astrocytes (S100) in cortex and dentate gyrus. Co-staining with NeuN and mCherry revealed that over 88% of the neurons imaged expressed mCherry driven by the different AAV-LAP variants in both cortex and dentate gyrus (Figure 6A1 - 6B4, 6E). Conversely, less than 4% of mCherry-positive oligodendrocytes were detected for all LAP variants (Figure 6C1 - 6D4, 6F). Moreover, we observed no co-labelling of mCherry with microglia (Iba1) and astrocyte (S100) markers for any of the AAV-LAP recombinants (Figure S1). Overall, these results demonstrate that in the context of systemic brain transduction with AAV-PHP.eB, LAP-mCherry expression is abundant in neurons but not in glial cells.

**Figure 6.**
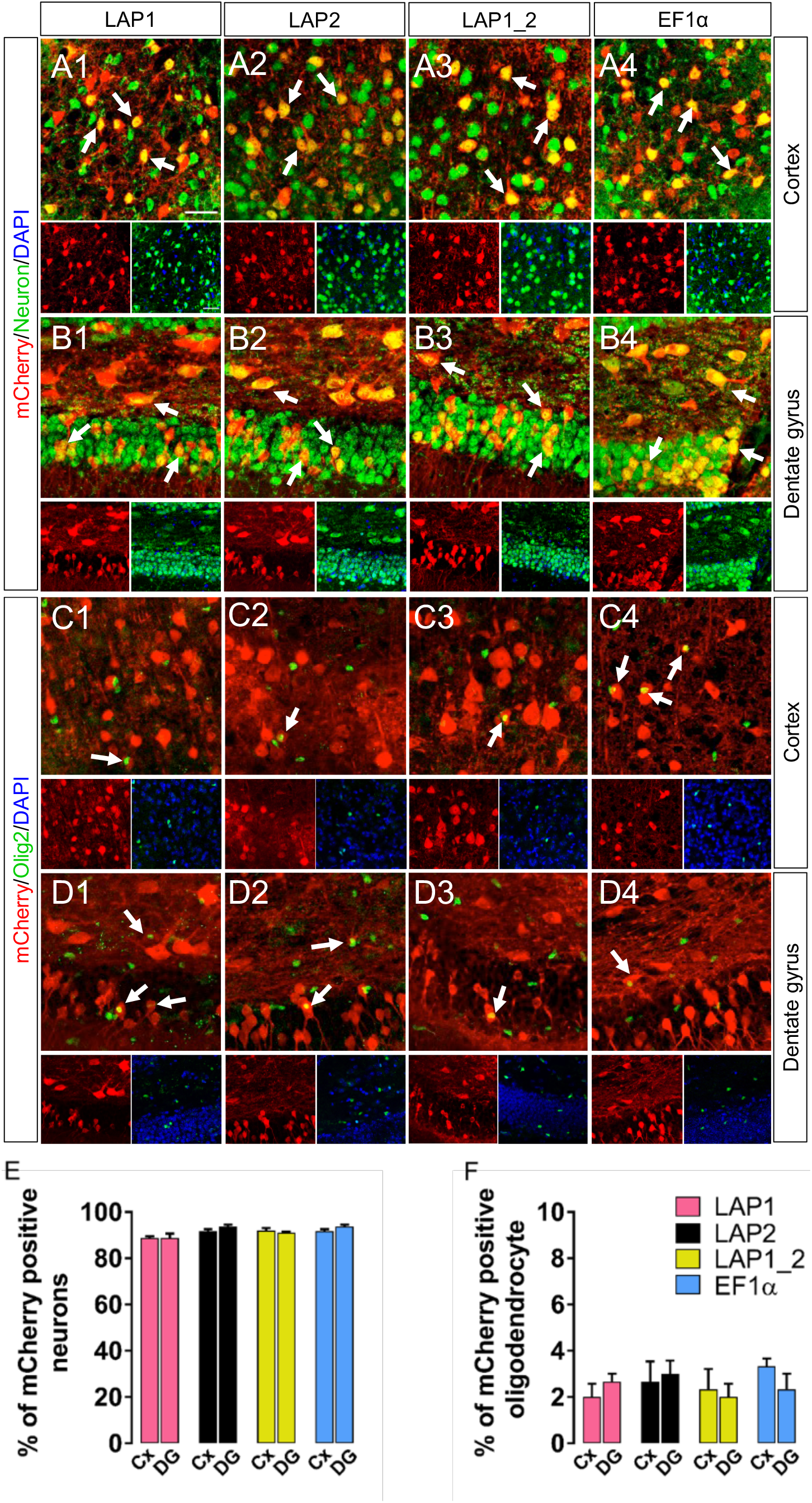
AAV-LAP transgene expression is predominantly expressed in neurons and not oligodendrocytes. (A, B) Representative confocal images of AAV-mediated mCherry expression (red) in neurons (green label for the pan-neuronal marker NeuN); (C, D) oligodendrocytes (green label for the oligodendrocyte marker Olig2), in both cortex and dentate gyrus at 30 dpi. Cells were counterstained with DAPI (blue). NeuN signal can localize with the neuronal cell nucleus as well as the cytoplasm, while the staining for Olig2 signal is mostly nuclear. Arrows depict co-labelling between the cell marker and mCherry. (A1, C1) AAV-LAP1 in cortex, (B1, D1) AAV-LAP1 in dentate gyrus; (A2, C2) AAV-LAP2 in cortex, (B2, D2) AAV-LAP2 in dentate gyrus; (A3, C3) AAV-LAP1_2 in cortex, (B3, D3) AAV-LAP1_2 in dentate gyrus and (A4, C4) AAV-EF1α in cortex, (B4, D4) AAV-EF1α in dentate gyrus. Scale bar = 100 μm. (E) Quantification of the percentage of AAV-mCherry labelled cells corresponding to neurons (NeuN-positive) or (F) oligodendrocytes (Olig2-positive) in cortex (Cx) or dentate gyrus (DG) for each promoter respectively. Images are stack confocal sections. Data are represented as mean ± SEM; n = 2 (six tissue sections were analyzed per animal).

### AAV-LAP constructs exhibit broad, stable and long-term transgene expression throughout the spinal cord

In addition to the brain, we evaluated AAV-LAP performance in spinal cord, where the serotype PHP.eB has shown widespread transduction of gray matter^7, 8^. We observed abundant native mCherry expression in both dorsal and ventral horns of the spinal cord at cervical, thoracic and lumbar levels 190 dpi (Figure 7A – 7D). Native mCherry RFI was similar for all promoters with no statistically significant differences (p < 0.05) (Figure 7E). However, the LAP2 and LAP1_2 recombinants showed the highest density of mCherry-positive cells per pixels^2^, followed by EF1α and LAP1 respectively; LAP2=LAP1_2>EF1α>LAP1 (Figure 7F). Therefore, all three PRV LAP variants effectively mediate pan-neuronal, long-term transgene expression in the spinal cord.

**Figure 7.**
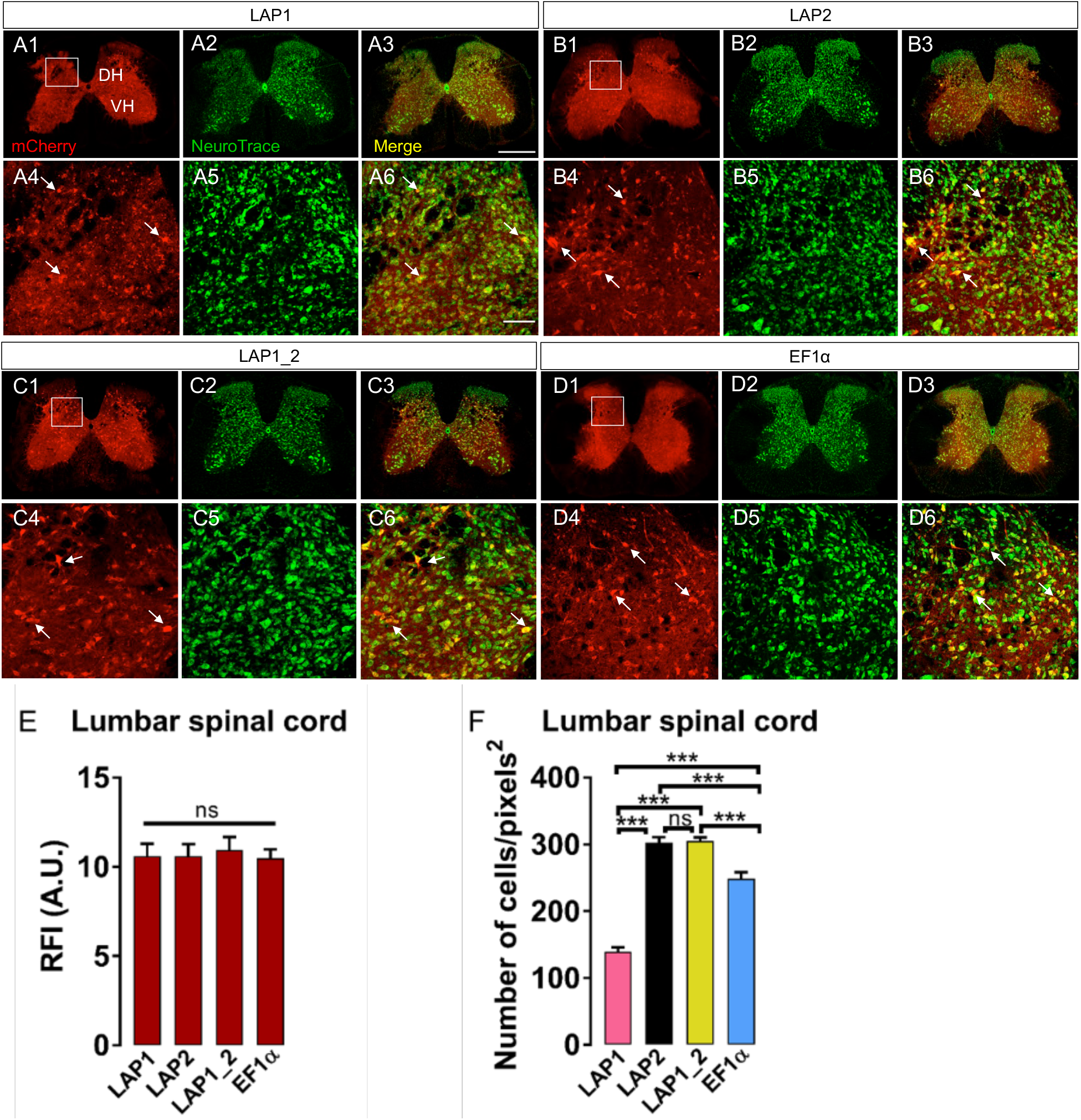
LAP drives widespread and long-term transgene expression in the spinal cord. Spinal cords (lumbar region) were sectioned in a transversal fashion at 20 μm. Representative 190 dpi confocal images from native AAV-mediated mCherry expression (red), pan-neuronal marker NeuroTrace (green), and merge signal (yellow) are shown for (A1 - A3) AAV-LAP1, (B1 - B3) AAV-LAP2, (C1 - C3) AAV-LAP1_2 and (D1 - D3) AAV-EF1α. DH, dorsal horn; VH, ventral horn. Higher magnification images of the DH are shown for (A4-A6) AAV-LAP1, (B4-B6) AAV-LAP2, (C4-C6) AAV-LAP1_2 and (D4-D6) AAV-EF1α respectively. Images are stack confocal sections. Scale bar = 1 mm and 100 μm.

## Discussion

Gene therapy has been used to restore gene function in specific target cells in neurological disorders^50^. Gene transfer by systemic vector delivery via peripheral vascular transduction can be difficult for efficient expression in a neuron-specific or pan-neuronal fashion in the CNS^51^. Recombinant AAV vectors are among the most efficient vehicles to achieve gene expression in the CNS^2,52^. Moreover, engineered AAV capsids have shown improved CNS transduction and enhanced capacity to cross the BBB with higher efficiency than naturally-occurring serotypes^6, 7, 53, 54^. Despite these advances, AAV gene therapy is hindered by the small payload size limit of 4.9 Kb for the AAV capsid^55^. For example, CNS therapies for Pompe disease ^56^ and Parkinson’s disease^57^ are based on delivery of relatively large genes such as GAA (2.9 Kb) and GDNF (2.5 Kb). For these and other similar cases, the use of small, non-repressible promoters are ideal replacements for larger promoters or even smaller CMV and hSyn promoters shown to be quickly repressed after delivery^14, 15^.

We identified three small pan-neuronal promoters isolated from the genome of the alphaherpesvirus PRV, showing efficient and long-term transgene expression in the mouse CNS after systemic AAV PHP.eB delivery. Our results demonstrate that these small PRV LAP variants can drive long-term expression of a reporter transgene (> 6 months) in brain and spinal cord. PRV LAP uniformly transduced neurons in the cortex, striatum, dentate gyrus and cerebellum. The distribution of mCherry-positive cells was not significantly different between LAP variants in the dentate gyrus, striatum and cerebellum. However, LAP2 transgene expression was significantly higher in cortex compared to LAP1 and LAP1_2. Our whole-brain screening assay demonstrated that the LAP2 variant of only 402 bp can drive stronger mCherry expression than the larger LAP1 and LAP1_2 sequences. Moreover, we detected abundant mCherry mRNA transcribed from LAP2 in every screened brain region at 190 dpi. These results demonstrate both efficient transcription and translation driven from the small PRV LAP2 in CNS after systemic AAV-LAP2 delivery in the absence of PRV infection.

Although the LAP1-mCherry cell density was significatively lower than that for LAP2, mCherry expression remained stable and long-lasting. Therefore, LAP1 might be useful in cases where low amounts of the therapeutic protein are needed (e.g. enzyme deficiencies) or for cross-correction to non-transduced cells. For example, for lysosomal enzyme deficiency^19^ and mucopolysaccharidosis VII diseases^58^, where the enzyme restored by AAV therapy can be secreted from the transduced cell and improve neighboring diseased cells.

AAV tropism is determined primarily by interactions between the capsid and specific receptors in susceptible and permissive cells ^4, 6^. Different AAV serotypes have different tissue tropism. However, the promoter sequence and other sequences included in the vector such as the inverted terminal repeat sequence (ITR), can have a substantial impact on tropism^59^. In addition, the promoter region transcribing the transgene is critically important to optimize the AAV vector’s performance. Efficient transgene expression either in a broad or cell-type specific fashion, requires binding and action of cell-derived TF to the promoter region^8, 60^. Changes in the neuronal environment such as aging or differentiation, also can alter the recruitment of cell-specific regulatory proteins and therefore gene expression in the CNS^61^. Our analysis of the PRV LAP sequence identified DPE in LAP2, which could control transgene expression onset, duration and cell-type specificity. Additionally, we identified four STAT1 motifs, that in HSV-1 LAP seem to regulate viral reactivation from latency^62^. Strikingly, we identified one of these STAT1 motifs in PRV LAP2 co-localizing with the TATA box and an Olig2 motif, a known multifaceted TF promoting neuronal and oligodendrocyte fates^63^. The proximity effects associated with these motifs and the transcriptional start site could explain the different levels of CNS transgene expression between LAP2, LAP1 and LAP1_2. However, further investigation of these regulatory elements is required to understand the fine-tuning control of PRV LAP2 activity. Additionally, we found one CTCF motif downstream of the LAP2 TATA box, which could have a role in the resistance to epigenetic silencing during latency, as shown for HSV-1^64, 65^. Indeed, Zimmerman and colleagues found that insertion of a CTCF motif downstream of the EF1α promoter increased transgene expression significantly compared to native EF1α and CMV promoters^66^. Interestingly, the insertion of a secondary CTCF motif downstream of the CMV TATA box, had no effect on luciferase reporter expression, presumably due to the redundant presence of a native CTCF motif^66^. Accordingly, gene expression is susceptible to changes depending of the genetic context and sequence-specific DNA binding proteins. The recruitment of specific TF from different host cells can modulate transgene transcription by the same mechanisms regulating resistance to inactivation during latency. Insulator elements like the CTCF-binding factor are independently regulated^67^ and can protect promoter regions from repression by heterochromatin, maintaining long-lasting transcription^64^.

Histological assessment of cell-specific transduction by colocalization of LAP-mCherry, glial and neuronal markers, revealed that LAP variants express more efficiently in neurons than glia, in cortex and dentate gyrus. LAP-mCherry positive cells colocalized predominantly with neuron-specific markers, and to a lesser extent with oligodendrocytes but not microglia or astrocytes. Based on these findings, we conclude that PRV LAP sequences have a pan-neuronal promoter profile in the CNS after PHP.eB systemic delivery. This activity has also been found for the HSV LAP due to the presence of a CRE motif upstream of the TATA box^68–70^. Transgenic mouse lines containing PRV LAP1 and LAP2 demonstrated that LAP are neuron-specific promoters in the absence of other viral proteins and that neuronal TF are sufficient to activate LAP *in vivo*^39^. Importantly, AAV-PHP.eB transduces neurons predominantly^7^, and the combination of this capsid variant with PRV LAP sequences exhibits a strong, long-lasting, pan-neuronal expression profile in the CNS. Future research should asses the transgene expression profile of PRV LAP and different AAV serotypes in tissue/organs other than the CNS and in animal models other than rodents. We predict that PRV LAP can be used not only in the context of recombinant viral vectors (AAV, adenovirus, lentivirus, herpesvirus), but also with non-viral gene delivery platforms. The natural host of PRV is the adult swine, but the virus has an extremely broad tropism and can infect some birds, fish and many types of mammals including some primates^71^. Moreover, human cells in culture are susceptible to PRV infection and there have been some reports of zoonotic infections^72^. Therefore, the PRV LAP sequence could be naturally optimized for gene therapy-applications requiring efficient and long-term transgene expression in several different mammals including humans.

In summary, we have demonstrated that PRV LAP promoter activity is independent of PRV infection and found that small AAV-PHP.eB-LAP variants express transgenes in a stable and pan-neuronal fashion in brain and spinal cord (CNS). Long-term transgene transcription and translation is paramount for effective and long-lasting single-dose gene therapy applications. Thus, PRV LAPs may be useful for the treatment of genetic CNS diseases after one-time viral-vector administration.

## Materials and Methods

### Construction of PRV LAP promoters

The PRV latency-associated transcript promoter (LAP) was PCR amplified from coordinates 95106-96007 of PRV Becker strain genome (GenBank: JF797219.1). The LAP1 region (498 bp), was amplified using primer pairs LAP1F (5’-GCA CGC GTA TCT CCG GAA AGA GGA AAT TGA -3’) and LAP1R (5’-GCG GAT CCT ATA TAC ACG ATG TGC ATC CAT AAT -3’). The LAP2 region (404 bp), was amplified using primer pairs LAP2F (5’-GCA CGC GTA TCC CCG GTC CGC GCT CCG CCC ACC CA -3’) and LAP2R (5’-GCG GAT CCG AGC TCC CTC TTC CTC GCC GCG GAC TGG -3’). LAP1_2 (902 bp) spanning the entire LAP region was amplified using LAP1F and LAP2R^31^. The 5’ and 3’ regions of these PCR sequences contained the MluI and BamHI restriction sites respectively, used for directional cloning into vector pAAV-EF1α-mCherry. The three AAV-LAP plasmids were constructed by double digestion of vector pAAV-EF1α-mCherry with MluI and BamHI followed by subcloning of the appropriate LAP fragment upstream of the mCherry reporter gene, flanked by AAV2 inverted terminal repeats (ITRs) and terminated with the SV40 polyA signal. pAAV-EF1α-mCherry was a gift from Karl Deisseroth (Addgene plasmid # 114470).

### Construction of AAV Vectors

All expression cassettes were packaged into AAV-PhP.eB capsids (gift from Daniela Gradinaru, Addgene plasmid # 103005) at the Princeton Neuroscience Institute Viral Core Facility and purified by iodixanol step gradient and column ultrafiltration as previously described^7, 73^. Capsid-protected viral genomes were measured by TaqMan qPCR and reported as genome copies per milliliter (GC/ml).

### Animals

Animal studies were performed following guidelines and protocols approved by the Institutional Animal Care and Use Committee of Princeton University (protocol # 1943-16 and 1047). Timed-pregnant Sprague-Dawley rats were obtained from Hilltop Labs Inc. (Scottsdale, PA). Adult (4 to 6-week-old) wild type C57BL/6J male mice were obtained from Jackson Laboratory (The Jackson Laboratory, Bar Harbor, ME). Mice had at least 48 hr of acclimation to the holding facility in the Princeton Neuroscience Institute vivarium before experimental procedures were performed.

### Primary superior cervical ganglia cell culture

SCG neurons from rat embryos (E17) were cultured in trichambers as previously described^74^. Briefly, SCG were dissociated with trypsin (2.5 mg/ml, Sigma-Aldrich, The Woodlands, TX) and plated on poly-O-Ornithine and laminin-coated dishes with media containing neurobasal media supplemented with 2% B-27, 100 ng/ml nerve growth factor (NGF), and 1% penicillin-streptomycin-glutamine (Thermo Fisher Scientific, Rockford, IL). Approximately two-thirds of a single ganglia were placed for S (soma) compartment of the trichamber. Three days post seeding, culture medium was treated with 0.1 mM cytosine-D-arabinofuranoside, Ara C (Sigma-Aldrich, The Woodlands, TX) for at least 2 days to eliminate dividing, nonneuronal cells. Culture media was replaced every 5 days, and neurons were incubated at 37 °C with 5% CO_2_.

### Retro-orbital sinus injection

Intravenous administration of AAV vectors was performed in mice by unilateral injection into the retro-orbital venous sinus^75^. Animals were anesthetized using ketamine (80 mg/kg)/xylazine (10 mg/kg) cocktail prior to the procedure. Once unresponsive, animals were placed in lateral recumbence for injection into the medial canthus. Injection volume was 100 μl containing a total of 4 × 10^11^ viral genomes administered with a 29G1/2 insulin syringe. Animals were placed on regulated heating pads and monitored until ambulant.

### Tissue processing and histological procedures

Mice were anesthetized with an overdose of ketamine (400 mg/kg)/xylazine (50 mg/kg) (i.p.) and perfused with 4% paraformaldehyde (PFA) at 30 and 190 dpi. Brain and spinal cord were post-fixed for 2 h in 4% PFA at RT (room temperature). After rinsing with phosphate buffered saline (PBS), brains were divided into two parts. Right hemispheres were used for iDISCO+ tissue clearing protocol (below). Left-brain hemispheres and spinal cords were serially incubated in 10% sucrose, 20% sucrose and 30% sucrose overnight at 4 °C. Tissue was placed in an embedding mold (Sigma-Aldrich, The Woodlands, TX) with OCT (tissue-Tek, Torrance, CA), froze in dry ice and stored at −80 °C until it was cryosectioned. Left hemispheres were sagittally sectioned at 50 μm using a Leica VT1200 vibratome for IHC and at 20 μm using a Leica CM3050 S cryostat for RNAscope. Spinal cords were transversally sliced at 20 um using a cryostat.

### Immunostaining

For immunohistochemistry, free-floating brain sections were washed with PBS and blocked for 1 h with 3% bovine serum albumin (BSA), 2% donkey serum and 0.5% Triton X-100 (Sigma-Aldrich, St. Louis, MO). Samples were incubated with primary overnight at 4 °C and secondary antibodies for 1 h at RT diluted in PBS containing 1% BSA, 1% donkey serum and 0.5% Triton X-100. Cell nuclei were counterstained with 0.5 μg/ml DAPI for 5 min (Thermo Fisher Scientific, Rockford, IL). The following primary antibodies were used: rabbit anti-RFP Rockland (1:1000; Limerick, PA), chicken anti-mCherry Abcam (1:500; Cambridge, MA), mouse anti-NeuN (1:500; Millipore Bioscience Research Reagents, Temecula, CA), rabbit anti-Olig2 (1:500; EMD Millipore, Temecula, CA), rabbit anti-Iba1 (1:1000; Wako Chemical, Richmond, VA) and rabbit anti-S100 (1:5000; Dako, Glostrup, Denmark). The following secondary antibodies were used: Alexa Fluor 488 donkey anti-rabbit IgG, Alexa Fluor 488 donkey anti-mouse IgG, Alexa Fluor 647 donkey anti-rabbit IgG, Alexa Fluor 647 donkey anti-chicken IgG (1:1000; Thermo Fisher Scientific, Rockford, IL). Spinal cord free-floating sections were stained with 1:300 dilution of NeuroTrace 500/525 green fluorescent Nissl stain (Molecular Probes, Eugene, OR) for 1 h. The sections were permeabilized with 0.1 % Triton X-100 in PBS at 10 min and washed first with PBS followed by PBS with 0.1% Triton X–100 for 10 min. Samples were incubated with 0.5 μg/ml DAPI for 5 min and then washed with PBS for 2 hours at RT. Fluoromount-G mount medium (Southern Biotech, Birmingham, AL) was applied to brain and spinal cord sections before mounting.

### Microscopy

Neuronal SCG cultures were imaged with a Nikon Ti-E inverted epifluorescence microscope (Nikon Instruments Inc, Tokyo, Japan), containing a Cool Snap ES2 camera (Photometrics, Tucson, AZ) and 4X objective. Tiled images of the entire S compartment were assembled with the Nikon NIS Elements software. To quantify AAV transduction efficacy in various brain regions, brain slices were imaged with a NanoZoomer S60 fluorescent microscope scanner (Hamamatsu, Hamamatsu, Japan). Brain slices were imaged with a Leica STP8000 confocal laser-scanning microscope (Leica Microsystems, Wetzlar, Germany) using 20X and 63X objectives, hybrid (HyD) detectors for sensitive detection, and over a 1024 × 1024 pixels area. Stacks of consecutive images were taken with a 20x objective and Z projections were reconstructed with ImageJ software to calculate corrected total cell fluorescence as previously reported^76^. Cells were selected drawing a region of interest (ROI) and normalized to background intensity from non-fluorescent cells. The calculation of corrected total cell fluorescence was measured as relative fluorescence intensity (RFI) considering area integrated, density and mean gray value of each cell.

### iDISCO+ tissue clearing

#### Permeabilization

Right brain hemispheres were used for iDISCO+ tissue clearing. Brain samples were fixed overnight in 4% PFA prior to tissue clearing as previously described^46^. Fixed samples were washed/dehydrated in 20, 40, 60, 80, 100% methanol/water solutions for 1 h each, followed by a 5% hydrogen peroxide/methanol overnight wash (Sigma-Aldrich, St. Louis, MO) and rehydration with a reverse gradient of methanol/water 100, 80, 60, 40, 20% for 1 h each. Finally, brains were washed with 0.2% Triton X-100 /PBS, followed by 20% DMSO (Thermo Fisher Scientific, Rockford, IL)/0.3M glycine (Sigma-Aldrich, St. Louis, MO)/0.2% Triton X-100/PBS at 37°C for 2 days.

#### Immunolabeling

Samples were incubated in a blocking solution of 10% DMSO/6% donkey serum (EMD Millipore, Temecula, CA)/0.2% Triton X-100/PBS at 37 °C for 2-3 days, followed by two 1 h/washes, in PBS/0.2% Tween-20 (Sigma-Aldrich, St. Louis, MO) with 10 μg/ml heparin (solution hereinafter referred to as PTwH, Sigma-Aldrich, St. Louis, MO). Brains were incubated with primary rabbit anti-RFP antibody (1:1000; Rockland, Limerick, PA) in 5% DMSO/3% donkey serum/PTwH at 37 °C for 7 days. Next, brains were washed with PTwH 5 times (wash intervals: 10 min, 15 min, 30 min, 1h, 2h) and incubated at 37 °C for 7 days with secondary Alexa Fluor 647 donkey anti-rabbit IgG (1:450; Thermo Fisher Scientific, Rockford, IL) in 3% donkey serum/PTwH and then washed in PTwH for 5 times.

#### Tissue clearing

Brains were sequentially dehydrated in 20, 40, 60, 80, 100% methanol/water for 1 h each step, followed by 2:1 dichloromethane (DCM; Sigma-Aldrich, St. Louis, MO)/methanol, and 100% DCM washes. Finally, samples were cleared with DBE (Sigma-Aldrich, St. Louis, MO) and stored in the dark at RT until imaged.

### Light-sheet microscopy and analysis of cleared tissue

After immunolabeling and clearing, brain volumes were acquired using a light-sheet Ultramicroscope II (LaVision Biotec, Bielefeld, Germany). Brain halves were glued in the horizontal orientation on a custom-designed 3D-printed holder^45, 46^ and submerged in DBE. Brains were imaged in the autofluorescent channel for registration purposes with a 488 nm laser diode excitation and a 525 nm maximum emission filter (FF01-525/39-25, Semrock, Rochester, NY), and at 640 nm excitation with a 680 nm maximum emission filter (FF01-680/42-25, Semrock) for cellular imaging of AAV infected cells (anti-RFP). Separate left- and right-sided illumination autofluorescent images were acquired every 10 micrometers (z-steps size) using a 0.017 excitation-sheet NA and 1.3x magnification. Left and right sided images were sigmoidally blended at the midline. Autofluorescent volumes were registered to the volumetric Allen brain atlas (2015) using affine and b-spline transformations, as described by Renier and colleagues^45, 46^. To account for movement during acquisition and different imaging parameters between channels, cell signal volumes were registered to autofluorescent volumes with an affine transform. Brain volumes were analyzed with our modified ClearMap software: “ClearMap_cluster” (github.com/PrincetonUniversity/clearmap_cluster), compatible with high performance computing clusters. Between five and ten 500 um volumes per brain region were analyzed for each animal to obtain the average local density measurements. For all analyzed samples, detected objects on brain edges and ventricles were eroded by 75 μm from the edge of the structure to minimize false positives.

### RNAscope in situ hybridization

Brain cryosections were mounted on superfrost plus adhesion slides (Thermo Fisher, Waltham, MA) and stored at −80 °C. RNA staining was performed using the RNAscope multiplex fluorescent reagent kit (Advanced Cell Diagnostics (ACD), Newark, CA) following the manufacturer’s protocol. The mCherry probe ACD# 431201-C2 was used. Slices were pretreated with protease IV for 30 min at 40 °C, followed by probe incubation for 2 h at 40 °C. Then, different amplifier solutions were performed for 30, 30 and 15 min at 40 °C. Signal was detected with TSA plus fluorescein system (Perkin Elmer, Waltham, MA). Incubation steps were done in the ACD HybEZ hybridization system. Slides were counterstained with DAPI for 30 s at RT. Finally, slides were mounted with VECTASHIELD Vibrance antifade mounting medium (Vector Laboratories, Burlingame, CA).

### Statistics

Statistical data analysis was performed using GraphPad Prism 7 software (GraphPad Software, Inc., La Jolla, CA). Two-tailed Student’s test was used to compare between two groups, and non-parametric one-way ANOVA test followed by a Bonferroni multiple comparison post-test to compare among multiple groups. A p value < 0.05 was considered to be statically significant. Data are represented as the mean with SEM.

## Acknowledgments

This work was supported by a Research Innovator Award from the Princeton Neuroscience Institute to EAE and LWE. We thank Professor Sam Wang for sharing consumables and lab equipment and Dr. Mahdi Kooshkbaghi for software assistance.

## Supplemental data

Supplementary Figure S1 and movie S2, S3, S4, S5 are available online.

## Supplemental Figures

**Figure S1.**
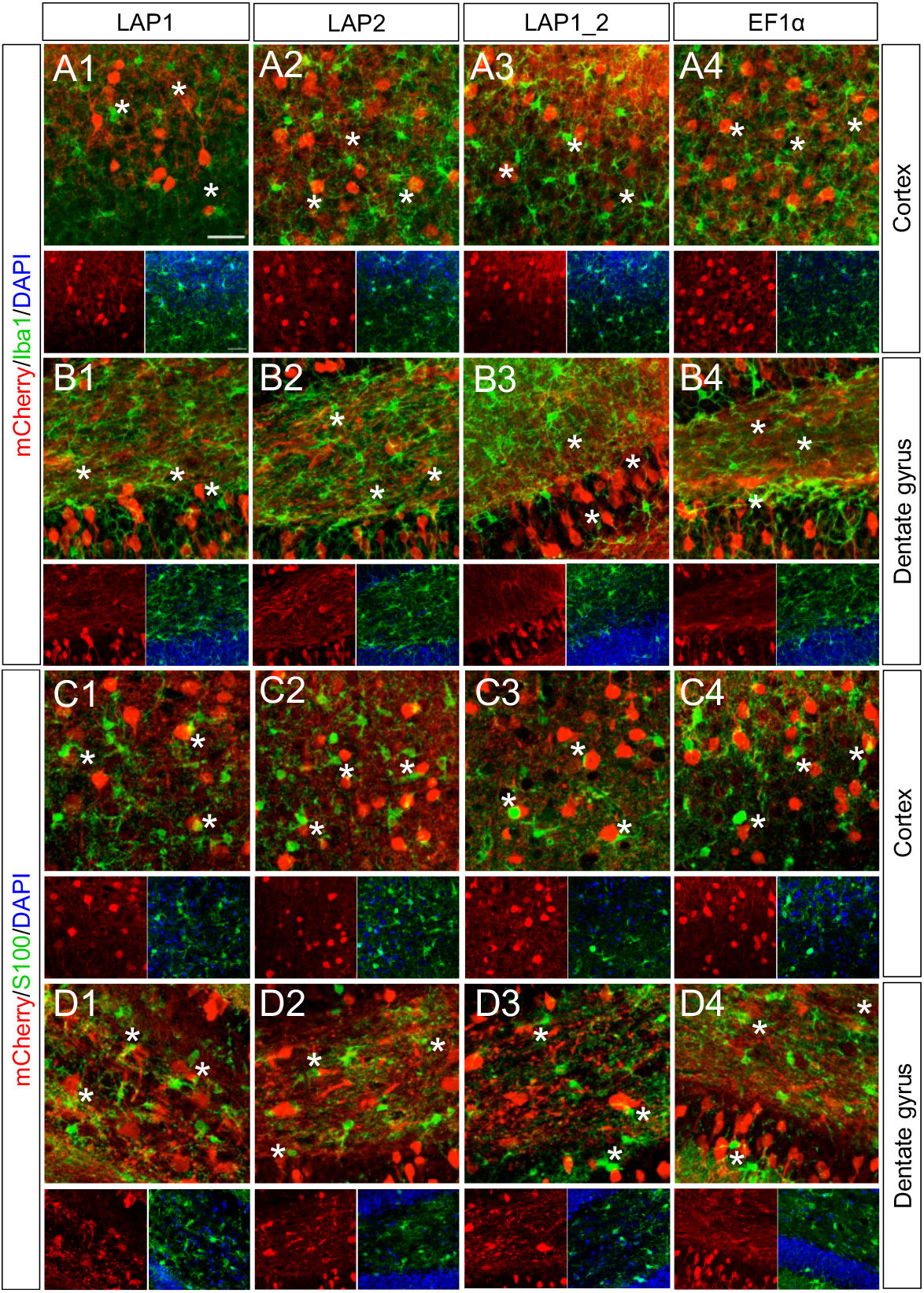
AAV-LAP transgene expression is not detected in microglia and astrocytes. (A, B) Representative confocal images of AAV-mediated mCherry expression (red) in microglia (green label for the glial marker Iba1); (C, D) astrocytes (green label for the astrocyte marker S100), in both cortex and dentate gyrus at 30 dpi. Absence of co-labelling of mCherry and astrocytes or microglia is indicated with stars (*). Cells were counterstained with DAPI (blue). (A1, C1) AAV-LAP1 in cortex, (B1, D1) AAV-LAP1 in dentate gyrus; (A2, C2) AAV-LAP2 in cortex, (B2, D2) AAV-LAP2 in dentate gyrus; (A3, C3) AAV-LAP1_2 in cortex, (B3, D3) AAV-LAP1_2 in dentate gyrus and (A4, C4) AAV-EF1α in cortex, (B4, D4) AAV-EF1α in dentate gyrus. Images are stack confocal sections. Scale bar = 100 μm.

